# Transcriptome Analysis of the Eggplant Fruits Overexpressing a Gene of Chlorogenic Acid Pathway

**DOI:** 10.1101/2020.10.09.332791

**Authors:** Prashant Kaushik

## Abstract

Chlorogenic acid is the primary phenolic acids present in Eggplant fruits. For the generation of chlorogenic acid in the eggplant hydroxycinnamoyl CoA-quinate transferase (SmHQT), is a central enzyme that catalyzes the reaction to the chlorogenic acid production. Although a precise function of SmHQT is not well defined in the eggplant fruit. In this study, the overexpression of SmHQT in the eggplant fruit’s flesh was studied using the agroinfiltration technique. Furthermore, to determine the differences at the genomic level RNA-seq analysis was performed, and its results showed that 415 genes of the phenylpropanoid pathway were upregulated in the transgenic fruit. Also, it was determined that the differentially expressed genes were dominantly related to phenylpropanoid pathway along with cell expansion, and cytokinin. The agroinfiltrated fruit exhibited more than twice the chlorogenic content in the normal fruit. Overall, the result provides new insight into the eggplant chlorogenic content increment at the molecular level and unseal the opportunities to design new strategies for the improvement of chlorogenic content as nutrition in eggplant.

## 1. Introduction

Phenolic acids are useful for human health as they have shown protection against diabetes, cancer and arthritis [1-3]. Eggplant flesh comprises of chlorogenic acid that makes up to 90% of the total phenolic acids present in the eggplant fruit flesh. For the production of chlorogenic acid, within the eggplant fruit flesh hydroxycinnamoyl CoA-quinate transferase (SmHQT), is the key enzyme which catalyzes the response on the chlorogenic acid output [4]. Therefore, improving the content of chlorogenic acid is one of the important breeding objectives for eggplant. Although cultivated eggplant has less phenolic acids compared to its wild species, the use of wild species in the eggplant breeding programs takes time to get rid of unwanted genes as the result of linkage drag [3]. Whereas with transgenic techniques, only the particular sequence is delivered in the genome that consequently expresses only the phenotypes associated with that gene [5]. Agroinfiltration is a technique that allows the transient gene expression in the plant tissues. Agroinfiltration approach for transient gene expression is well-tried in many fleshy fruits like tomato, strawberry, melon, and cucumber [6]. Thus, the objectives of this specific study had been studying the eggplant fruits flesh transiently overexpressing the SmHQT gene. Also, in our transgene cassette, we co-expressed the P19 gene of tomato bushy stunt virus as it prevents the post-transcriptional gene silencing (PTGS). Furthermore, the RNA sequencing (RNA Seq) analysis-based gene expression of agroinfiltrated fruits was also compared to regular fruits.

## 2. Materials and Methods

### 2.1. Plant Material, SmHQT Gene Construct and Agroinfiltration

Seeds of green and long cylinder fruit-bearing eggplant variety Arka Shrish were grown with a mixture of turf soil and perlite (2:1) under a greenhouse with natural light at 20–25 °C. The gene construct and agroinfiltration procedure followed is defined elsewhere [7]. Briefly, genomic DNA was extracted from the eggplant fruit and amplified for the SmHQT gene together with its native promoter. The optimized gene was cloned in the cloning vector (pUC57 vector; Addgene, Spain) at *Hind*III/*Bam*HI sites. The pUC57+p19 clone was restriction digested (*Hind*III/*Bam*HI); the released gene was later cloned into pBIN19 at (*Hind*III/*Bam*HI). A 2ml syringe with a needle was used to inject the agro culture in Eggplant fruits at 10-15 spots. After that, the plants were left to grow for 3 to 10 days after infiltration (DAI). Fruit samples were harvested from the infiltrated plants at 3 DAI. After that, fruits were harvested and stored in −80°C for further studies.

### 2.2 Validation and Quantification

The positive expression was compared in the agroinfiltrated fruit as compared to the control by X-Gluc staining based on the GUS gene. Briefly, 1mM X-Gluc was prepared according to standard protocol, and the fruit sections were stained for 30mins 200 at 37°C and visualized under a light microscope (LYZER LT-1610X). Also, the quantification of the chlorogenic acid content (g/kg of fresh weight) was performed based on the detailed protocol defined elsewhere using the high-performance liquid chromatography (HPLC) method with a 1220 Infinity LC System (Agilent Technologies, Santa Clara, CA, USA). The analysis part was carried out by the OpenLAB CDS ChemStation Edition program (Agilent Technologies) [8,9].

### 2.2. RNA-sequencing and Data Analysis

The RNA was extracted from the three agroinfiltrated fruits of and control fruit for RNA isolation whole fruit were taken after peeling the RNA was extracted with the help of Qiagen RNeasy Plant Mini Kit (Qiagen, Valencia, CA) with precisely following manufactures instruction protocol. The RNA integrity was determined using a bioanalyzer (Agilent Technologies 2100), and samples with an RNA Integrity Number (RIN) value greater than or equal to seven were processed for further use. After that, the RNA from control and the transgenic plant was pooled in equimolar concentrations as separate pools for library preparation. RNA Library Prep Kit v2 from Illumina® (Illumina Inc., San Diego, CA, USA) was used for the preparation of the two separate libraries, and the libraries were quantification with Qubit Fluorometer (Qubit™ dsDNA HS Assay Kit, Life Technologies Corporation, Carlsbad, California, USA). Illumina HiSeq 2500 (2 × 150 bp) platform (Illumina, Dedham, MA, USA) was used for the sequencing of the libraries. The raw data of RNA-seq is submitted to the BioProject with the project id PRJNA531188.

### 2.3. Quality Control and Transcriptome Assembly

The reads were quality filtered, utilizing Cutadapt (version 1.8.1). After adapter removal, the low-quality bases (q &lt;20) were eliminated by Trimmomatic software (version 0.39). The filtered reads of each of the samples had been merged for a de novo transcriptome assembly using Trinity assembler version (v2.11.0) that was enhanced for Illumina paired-end data and configurable for most computing capacities depending on the default parameter [10]. Then transcripts from the assembly were clustered going with the CD-HIT-EST application with criteria sequence identity threshold as 0.9 plus length difference cutoff as 0.9 to get the set of non-redundant transcripts (unigenes).

### 2.4. Differential Gene Expression Analysis

The unigenes were analyzed further for annotation and expression profiling. The expression of the unigenes were estimated by RSEM [11] and Bowtie 2 [12] software both are already configured with Trinity programme. After this the unigene count values were used as input to DESeq package for differential gene expression analysis. Significant expression of the transcripts was selected based on the criteria P<0.05 [13].

### 2.5. Functional Characterization

The unigene functions were annotated by subjecting for blastx against databases such as NCBI NR, PRIAM and PFAM in the FunctionAnnotator platform (E-value < 1e-5) [14]. The Gene Ontology (GO) terms were assigned for the unigenes by Blast2Go suit included in the FunctionAnnotator platform [15]. KAAS annotation server was used to perform the pathway analysis. For pathway analysis plant pathways were considered as references. The heatmap of top upregulated and downregulated genes were drawn using R package [16].

## 3. Results and Discussion

### 3.1. Argoinfiltration Assay and HPLC

Similarly, transient expression was also done with the native promoter. Transgenic plants obtained by Agrobacterium tumefaciens-mediated transformation of fruit were sent to the greenhouse; 3 DAI. Among the twenty fruits overexpressing SmHQT, fruits showed phenotypic variations in comparison to normal plants. The overexpression was confirmed by the X-Gluc staining, as shown in Figure 1. Furthermore, the chlorogenic acid content in the agroinfiltrated fruit flesh was more than twice than that of normal fruit (Figure 2).

**Figure 1.**
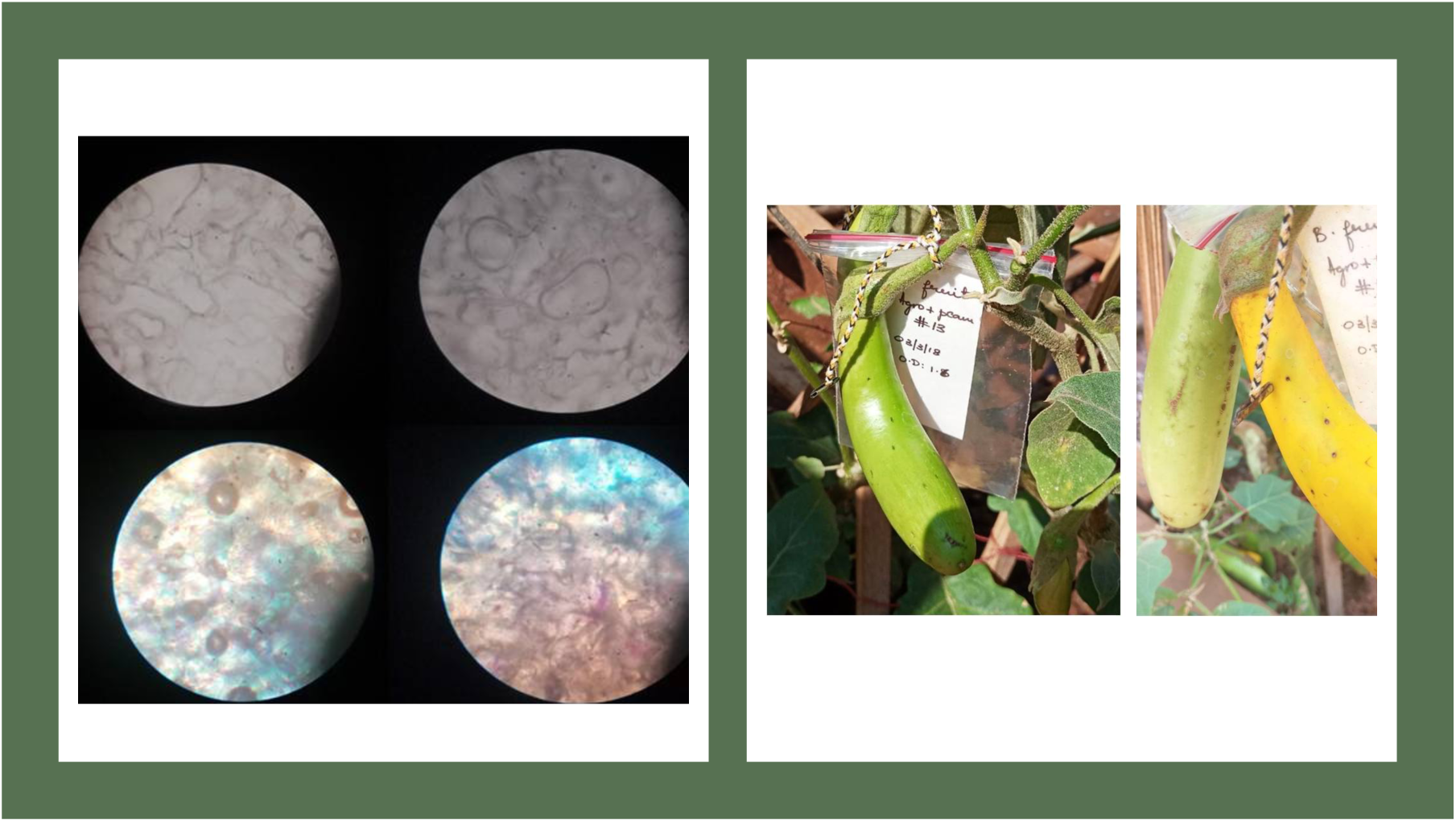
X-Gluc staining (on left side) of control fruit samples (above) vs agroninfiltrated (below). The fruits samples (on the right side) at 3 Days after agroinfiltration (yellow colored) vs control (green colored).

**Figure 2.**
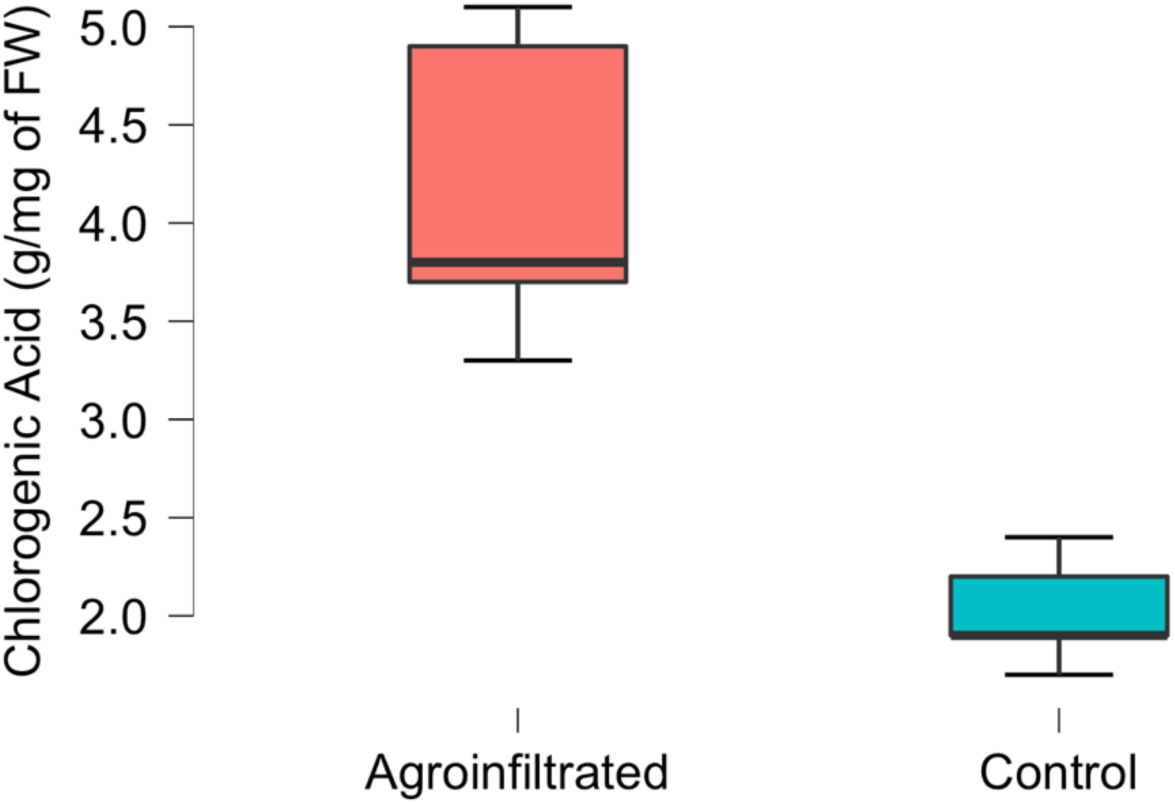
Chlorogenic acid Acid (g/mg of FW) in agroinfiltrated vs control fruit samples.

### 3.2. De Novo Assembly

The samples of SmHQT OE genes in eggplant were selected after agroinfiltration period of 3 days of infection to determine differentially expressed genes (DEGs). A total of 1372 short reads were determined from cDNA libraries of OE SmHQT in fruits.

### 3.6. DEGs Analysis and Gene Ontology (GO) Enrichment Analysis

The expression between controlled and agroinfilrated OE gene sample was compared in fruits after three days of infection in eggplant fruits. To that effect, an increased quantity of DEGs was determined in OE genotypes compared to the control (Figure 3). A total of 3166 and out of total 134 were combinedly expressed in both conditions meanwhile 1238, and 1794 are unique DEGs were significantly depicted in agroinfiltrated and control, respectively (Figure 3). Correspondingly, we determined around 694 DEGs were downregulated in response to OE, but we cannot say none of the genes were silenced in control. Furthermore, downregulation of 169 DEGs in reaction to OE was noticed and both control of chlorogenic biosynthesis pathway. GO analysis pointed out that the different functional groups (Figure 4). The groups like catalytic activity, transporter activity, as well as transcription regulatory activity (Figure 4). Likewise, a considerable upregulation selection of DEGs was noticed in the cellular component category in OE genotypes as compared to control fruit sample (Figure 4).

**Figure 3.**
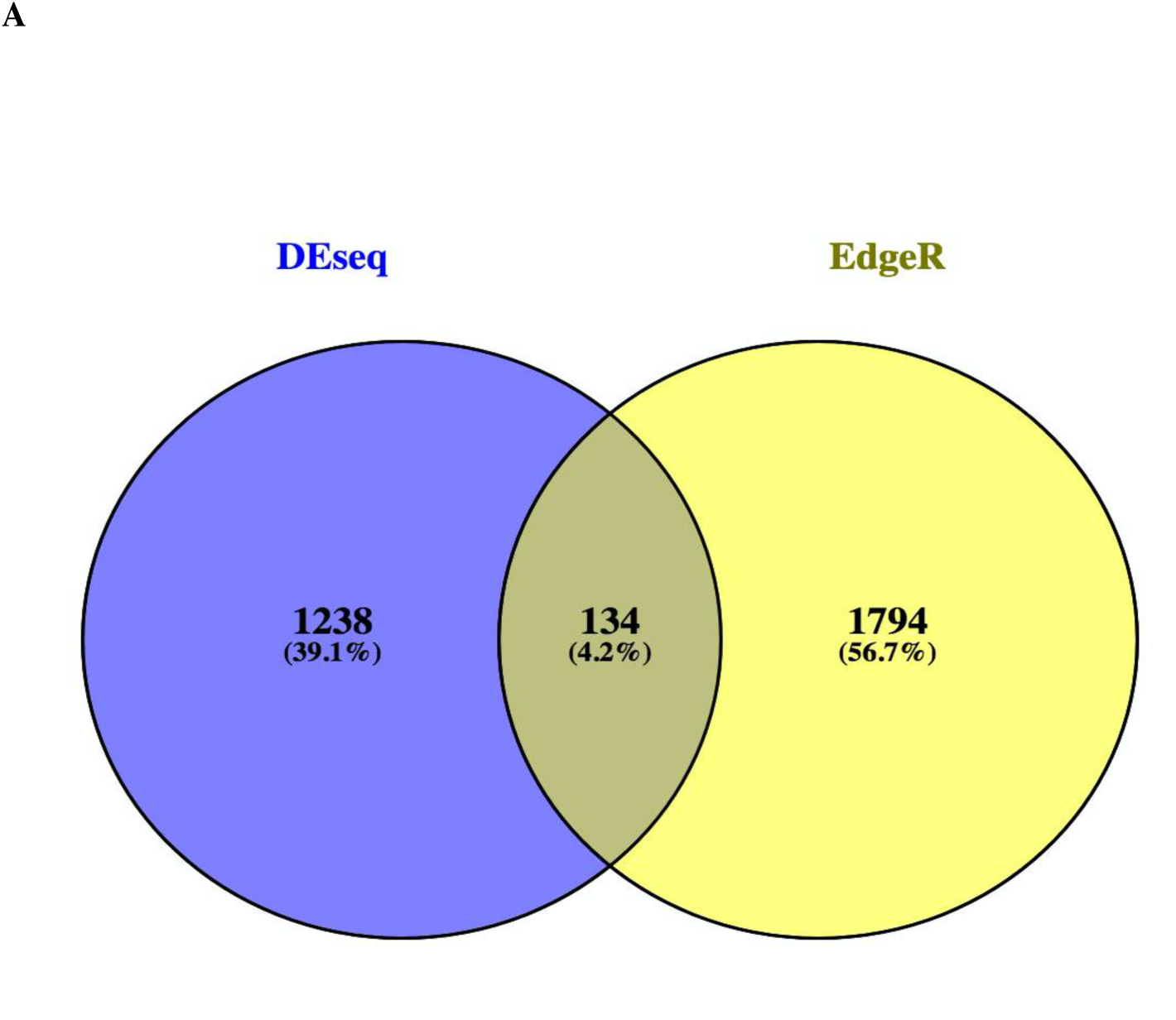
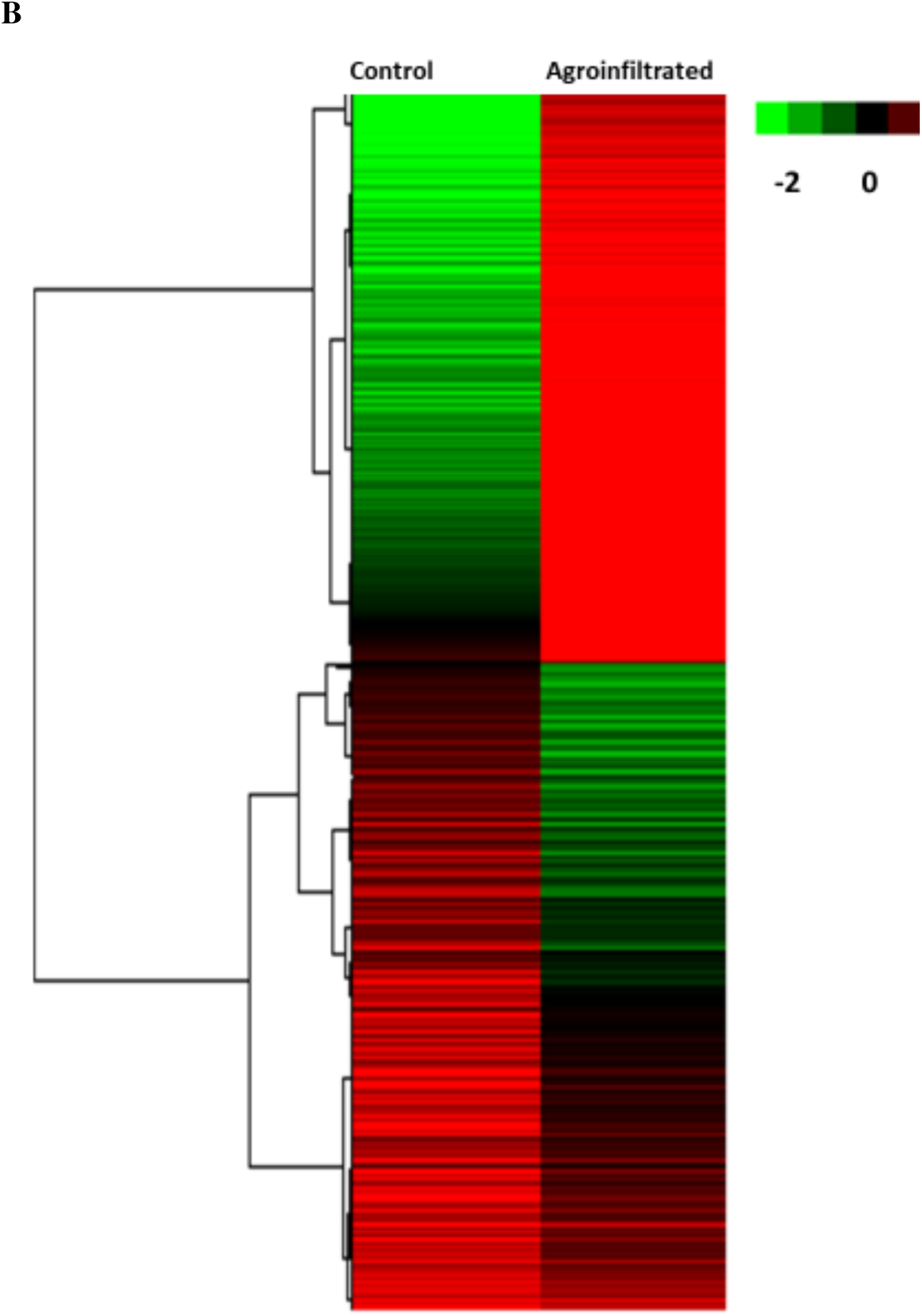
Venn diagrams indicating the number of DEG genes showing expression in and among control and agroinfiltrated fruits. The DEG values were determined from the NGS sequencing data (A).Heat map of transcripts in control vs agroinfiltrated sample of the eggplant fruit (B).

**Figure 4.**
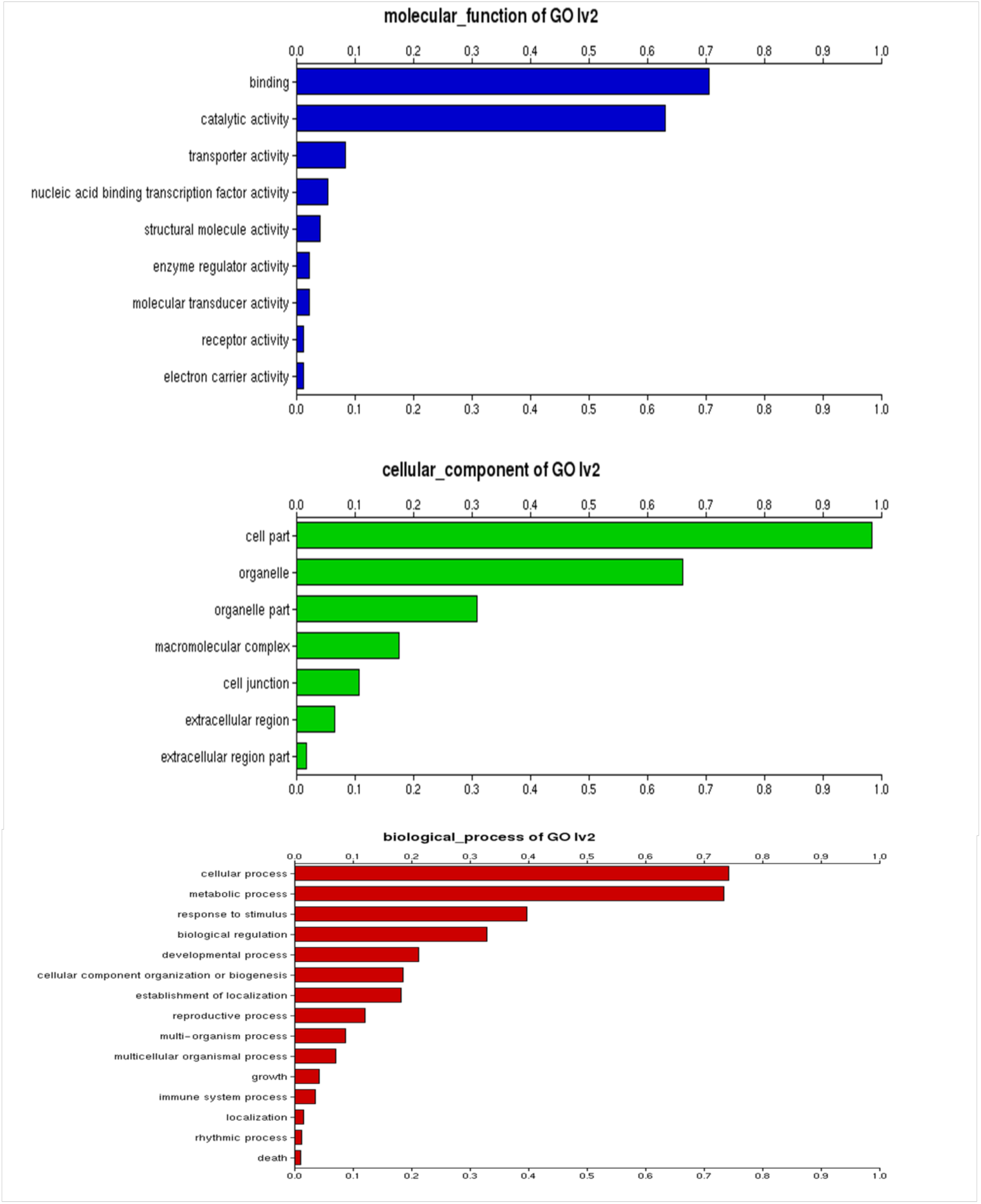
Gene Ontology (GO) terms identified in the eggplant fruit tissues.

Agroinfiltration is a compatible method for transiently expressing or silencing a gene. This method could be used to make vaccines and enzymes for industrial usage as an alternative to the conventional method of Agrobacterium transformation. Agroinfiltration studies in the past have supplied useful insights into many vital processes, like promoter function, gene expression, subcellular localization of the protein and metabolism. Agroinfiltration, when linked to RNA-seq, becomes a potent tool that can be to plant species where agroinfiltration protocols are optimized (strawberry, tomato and melon) [17]. For a great deal of plant life, including eggplant producing transgenic with a stable gene expression could be time-consuming and challenging [18].

## 4. Conclusion

The very first report and protocol on the impact of SmHQT overexpression on gene expression in eggplant for enhancement of chlorogenic content. Functional assessment of SmHQT gene suggested that overexpressing the SmHQT exhibited a higher content of chlorogenic acid, hinting its function as a good regulator of phenolic acids. Nevertheless, the possibility is present that the eggplant during overexpression phenotype is actually in part regulated by extra interacting partners genes determined in our RNA seq data

## Acknowledgements

The author is also thankful to the anonymous reviewers for their careful reading of the manuscript and for providing insightful suggestions.

